# Brain Vasculature Accumulates Tau and Is Spatially Related to Tau Tangle Pathology in Alzheimer’s Disease

**DOI:** 10.1101/2024.01.27.577088

**Authors:** Zachary Hoglund, Nancy Ruiz-Uribe, Eric del Sastre, Benjamin Woost, Joshua Bailey, Bradley T. Hyman, Theodore Zwang, Rachel E. Bennett

**Affiliations:** Department of Neurology, Massachusetts General Hospital, Charlestown, MA, USA; Harvard Medical School, Boston, MA, USA

**Keywords:** Alzheimer’s disease, tau, neurofibrillary tangles, blood vessels, cerebral vasculature, cerebral amyloid angiopathy

## Abstract

Insoluble pathogenic proteins accumulate along blood vessels in conditions of cerebral amyloid angiopathy (CAA), exerting a toxic effect on vascular cells and impacting cerebral homeostasis. In this work we provide new evidence from three-dimensional human brain histology that tau protein, the main component of neurofibrillary tangles, can similarly accumulate along brain vascular segments. We quantitatively assessed n=6 Alzheimer’s disease (AD), and n=6 normal aging control brains and saw that tau-positive blood vessel segments were present in all AD cases. Tau-positive vessels are enriched for tau at levels higher than the surrounding tissue and appear to affect arterioles across cortical layers (I-V). Further, vessels isolated from these AD tissues were enriched for N-terminal tau and tau phosphorylated at T181 and T217. Importantly, tau-positive vessels are associated with local areas of increased tau neurofibrillary tangles. This suggests that accumulation of tau around blood vessels may reflect a local clearance failure. In sum, these data indicate tau, like amyloid beta, accumulates along blood vessels and may exert a significant influence on vasculature in the setting of AD.

## Introduction

The formation of tau-containing neurofibrillary tangles (NFTs) is closely associated with the severity and progression of Alzheimer’s disease (AD)^1, 2^. Because of this close relationship, it is important to investigate the mechanisms by which these pathological aggregates of tau protein form, and why certain neurons are more vulnerable to tangle formation than others. On the molecular scale, NFTs form when tau protein is post-translationally modified, notably by phosphorylation at multiple sites in the protein’s structure, increasing its propensity to self-aggregate. In AD, aggregates develop in distinct regional patterns, including with varying density between cortical layers^3, 4^. These observations indicate that neuroarchitecture plays a significant role in the progression of tau pathology. In addition to intracellular accumulation in neurons, tau is also secreted into the extracellular space and can be detected in cerebrospinal fluid and blood.

Given that that tau is present extracellularly, we hypothesized that one key mechanism that may influence the local accumulation of tau pathology could be vascular brain clearance pathways^5^. Peri- and para-vascular clearance pathways serve as important routes for the removal of brain solutes, linking the interstitial (ISF) and cerebrospinal fluid (CSF) compartments, while trans-vascular clearance may directly transport other molecules from brain to the blood. Reduced export of protein wastes along these pathways is believed to lead to the accumulation of toxic and aggregation-prone species of AD-related proteins^6–9^. This is most clearly exemplified by cerebral amyloid angiopathy (CAA), a common AD co-pathology in which insoluble amyloid beta accumulates in the basement membrane and smooth muscle cells of leptomeningeal and penetrating arterioles in the brain. In a recent study, Harrison *et al* showed that globally perturbing the CSF-ISF flow accelerated tau deposition in the brains of a mouse model^6^. This work demonstrated that this system is important not just for amyloid beta but also for tau protein clearance; however, it did not examine individual vessels to understand their direct contribution to pathology. In related work, our group observed that isolated vasculature from both tauopathy mice and human AD brains contains high levels of bioactive tau capable of seeding new aggregates^10^. Together, this suggests that impaired clearance of these bioactive tau species could result in vasculature becoming important reservoirs for tau protein.

In this study, we identified the presence of tau immunoreactivity along cerebral brain vessels in AD patients and sought to quantify the relationship between tau pathology and brain vasculature at smaller, single-cell spatial scales. We predicted that if impaired perivascular clearance is related to formation of tau tangles, increased phosphorylated tau species would be present along blood vessels in the brain and tangles would exhibit close spatial relationships with vasculature compared to non-tangle bearing neurons. However, one challenge of investigating protein distribution in the brain is imaging structures at high resolution in large tissue volumes. Advances in tissue clearing and multiplexed antibody staining have addressed this gap and enabled us to quantify the distribution of tau pathology and determine its spatial relationship to vasculature in the AD brain with single-cell resolution^11, 12^. This allowed us to quantify, in three dimensions, the proximity of phosphorylated tau and NFTs to individually segmented blood vessels. In addition, we conducted protein assays to determine the presence of bioactive tau species in vasculature. Surprisingly, these experiments uncovered new evidence that, like CAA, tau accumulates along vascular segments in the AD brain. Additionally, NFT density positively correlated with the amount of tau accumulated along vascular segments, indicating that tau accumulation along vasculature is associated with tangle formation in Alzheimer’s disease.

## Materials and Methods

### Human tissues

Fresh frozen human tissue samples of the inferior temporal gyrus were provided by the Massachusetts Alzheimer’s Disease Research Center (ADRC) with approval from the Mass General Brigham IRB (1999P009556) and with informed consent of patients or their relatives. In total, 7 human participants with Alzheimer’s disease and 6 controls were selected from the Massachusetts Alzheimer’s Disease Research Center. Sex, age at death, Braak staging, post-mortem interval and comorbidities are listed in **Table 1**.

**Table 1:** Human tissues used in this study. Details include sex (M = male; F = female), age at death, Braak stage, comorbidities (CVD = cerebrovascular disease; LBD = Lewy Body Dementia; CAA = Cerebral Amyloid Angiopathy), and post-mortem interval (PMI; hours). None of the inferior temporal gyrus areas examined contained overt vascular lesions. Sample AD 7 was used for SMA labeling (Fig. 2F) while the others were used for quantitative assessments.

| Sample Number | Sex | Braak Stage | Age at Death | Comorbidities | PMI |
| --- | --- | --- | --- | --- | --- |
| AD 1 | M | V | 74 | CVD | 10 |
| AD 2 | M | VI | 75 | CVD | 4 |
| AD 3 | M | III | ≥90 | CVD | 14 |
| AD 4 | F | VI | 66 | CVD | 14 |
| AD 5 | F | VI | ≥90 | Arteriolosclerosis | 5 |
| AD 6 | F | VI | 70 | CVD | 24 |
| AD 7 | M | V | 52 | LBD, CAA, CVD | 28 |
| Control 1 | F | I | 68 | CVD | 20 |
| Control 2 | M | 0 | 62 | CVD | 17 |
| Control 3 | M | 0 | 73 | CVD | 14 |
| Control 4 | M | 0 | 63 | CVD | 18 |
| Control 5 | F | I | 64 | CVD | 18 |
| Control 6 | F | 0 | 56 | Acute Hypoxia | 8 |

## Protocol for assaying tau extracted from blood vessel homogenates

### Isolation of blood vessels

Blood vessels were isolated from 200-300 mg of frozen mice and human tissue. Brains were minced in 2 mm sections using a razor blade in ice-cold B1 buffer (Hanks Balanced Salt Solution with 10mM HEPES, pH 7; Thermo Fischer Scientific). Then samples were manually homogenized using a Dounce homogenizer with 12 strokes. Homogenate was then transferred into a conical tube filled with 20 mL of B1 buffer and centrifuged at 2,000 g for 10 minutes at 4 °C. Supernatant was discarded and pellet vigorously resuspended during 1 min in 20 mL of B2 buffer (B1 buffer with 18 % dextran, Sigma-Aldrich) to remove myelin. Samples were centrifuged 4,400g for 15 min at 4 °C. The myelin layer was carefully detached, and the pellet was resuspended in 1 mL of B3 buffer (B1 buffer with 1 % Bovine Serum Albumin, BSA, Sigma-Aldrich). Afterwards, homogenate was filtered through a 20 µm mesh (Millipore) previously equilibrated with 5 mL of ice-cold B3 solution. Brain blood vessels were rinsed with 15 mL of ice-cold B3 solution and then the blood vessels detached from the filters by immersing them in 20 mL of B3 ice-cold solution. Vessels were centrifuged at 2,000g for 5 min at 4 °C. Finally, the pellet was resuspended in 1 mL of ice-cold B1 solution and again centrifuged at 2,000g for 5 min at 4 °C and the supernatant was discarded. Vessel-containing pellets were stored at – 80 °C.

### Protein assays

Protein was extracted from human and mice brain blood vessels homogenates, which were sonicated at 20 % amplitude in 10 strokes in PBS supplemented with protease and phosphatase inhibitors (cOmplete Mini and PhosSTOP EASYpack; Roche). Then, samples were centrifuged at 3,000 g for 5 min at 4 °C and supernatant discarded. Proteins were analyzed following a capillary based electrophoresis instrument (SimpleWes, Biotechne). Three mg of protein were used per sample. Protein separation and detection were performed by capillary electrophoresis, binding of antibodies and HRP conjugated secondaries were done in SimpleWes machine. Antibodies used were phospho-T181 (mouse 1:50, MN10050, Invitrogen), phospho-S202 (rabbit 1:25, 39357S, Cell Signaling), phospho-T217 (rabbit 1:25, 44-744, Invitrogen), phospho-T231 (rabbit 1:50, #44-746, Invitrogen), Tau13 (mouse 1:50, 835201, Biolegend), Tau46 (mouse 1:50, 4019S, Cell Signaling) and total tau (rabbit 1:50, A0024, DAKO). Specific SimpleWes secondary antibodies HRP conjugated were acquired from the manufacturer (Biotechne). Protein quantification was analyzed in Fiji (https://doi.org/10.1038/nmeth.2019). The total intensity of signal in each lane was measured and normalized to the average of the three control samples.

## Protocol for Tissue Clearing and Imaging

*Tissue slicing.* Brain samples were placed in 4% paraformaldehyde (Thermo Fisher Scientific, cat No. 50980487) for 24 hours at 4°C. Tissue was then rinsed three times with 50 ml phosphate-buffered saline (PBS) for 10 minutes each, then placed in fresh PBS overnight at 4°C and rinsed with fresh PBS. Fixed tissue underwent three rinsing cycles in 10-minute increments using 50 ml of PBS, then were placed in fresh PBS overnight at 4°C. In preparation for tissue slicing, tissue was transferred to individual 35 mm Petri dishes and embedded in a gel block by pouring warm 4% agarose gel solution in PBS (4 g/100 ml) (Promega, cat No. V3121) over the tissue. The gel was then cooled to solidify and cut into a block to provide rigidity for cutting even slices. The tissue was secured on Vibratome (Leica Biosystems, VT1000 S Vibrating Blade Microtome) slide via super gluing the bottom of the agarose block. The vibratome was then used to slice 0.5–1 mm thick sections of tissue. Each slice was then removed from the agarose through gentle manipulation with blunt forceps or paintbrushes and were then placed in crosslinking solution, described below.

### Delipidation

Tissue was then placed into sodium dodecyl sulphate (Sigma-Aldrich, cat No. L3771) 28.83 g/500 ml PBS-clearing solution supplemented with sodium borate (Sigma-Aldrich, cat No. S9640) on shaker at 100 rpm and 37C for ∼3 days. After delipidation, the brain slices were rinsed with 50 ml PBS five times over 24 hours.

### Immunohistochemistry

Each brain slice was placed in a 2 ml Eppendorf tube that could hold the slice so its large, flat sides could be exposed to solution. PBST (PBS with 0.2% Triton X-100, Thermo Fisher Scientific) was added to just cover the top of the samples (∼500 μl). Tissue was heated to 50°C for 1 hour in PBST and then cooled to room temperature prior to incubation with antibodies. The following conjugated antibodies were then added to the solution containing each tissue slice: phospho-tau Ser202, Thr205 (AT8, 1.6:500, Thermo Fisher, cat No. MN1020) conjugated to Alexa Fluor 647 (Thermo Fisher, cat No. A37573), HuD Antibody E-1 (1.6:500, Santa Cruz Biotechnology, cat No. sc-28299) conjugated to Alexa Fluor 555 (Thermo Fisher, cat No. A37571), Glut1 antibody conjugated to Alexa Fluor 488 (EMD Millipore, 07-1401-AF488) and 4′,6-diamidino-2-phenylindole dihydrochloride (DAPI, 1.6:500, Sigma-Aldrich, cat No. 10236276001). Tissue was incubated with primary antibodies for one week at 4°C with gentle shaking. Following incubation, tissue was washed in fresh PBST 3 × 10 min and set on shaker for one week at 4°C with gentle shaking.

### Refractive index matching

After immunohistochemical staining, the samples were incubated with 80% glycerol, 20% deionized water for 24 hours at room temperature with gentle shaking. Samples were then placed on a glass microscope slide with a 3D-printed ring that allows the tissue to remain in a pool of glycerol during imaging. The ring was 3D-printed to match the thickness of the tissue (Formlabs) so a glass coverslip could be placed on top and seal the tissue in the glycerol.

### Imaging

The tissue was imaged using Olympus Inverted Confocal FV3000 with a 10 × air objective, and multi-region images were stitched together using the microscope software (Fluoview FV31S-SW, Version 2.5.1.228). Additional higher resolution images were collected by placing the tissue in a bath of 80% glycerol in a Petri dish and imaged with using a 20 × immersion objective (Zeiss Clr Plan-Neofluar 20x/1.0 Corr) with an inverted Zeiss 980 confocal microscope. Image Z-stacks were then reconstructed and visualized using Imaris microscopy image analysis software.

## Protocol for Segmentation and Quantification of Pathology

### Analysis with Ilastik (1.4.0)

Imaging data for tau and HuD were converted to HDF5 format using Ilastik’s ImageJ plugin^13^. The staining was then individually segmented for each image using Ilastik’s pixel classifier workflow. In short, a paintbrush was used to draw over the signal and background to help train the classifier on how to segment each image. All images were then processed through the trained pixel classifier, and probability maps were exported as HDF5 formatted images. Pixel probability maps and raw data were loaded into Ilastik’s object classification workflow and used to train object classifiers for each image. Tau object classifiers were trained by manually classifying objects as noise or tangles, and HuD object classifiers were trained by manually classifying objects as noise or neurons. Data was exported as object identities and spreadsheets with information about the objects’ classification and characteristics, which were then loaded into MATLAB (r2023b) code to match objects from each channel with colocalized objects.

### Separating objects into cortical layers

Imaris surface generation was used to draw regions around each cortical layer on individual imaging planes within the Z-stack, which was then merged into distinct volumes that contain each cortical layer. These volumes were then used to generate a new channel by masking the pixels contained within each volume and setting them equal to the cortical layer (i.e. pixels in layer 1 = 1, pixels in layer 2 = 2, etc.) and pixels not within a clearly defined layer equal to zero. This channel was then exported as a single multipage tiff stack, which could be loaded into our MATLAB code to identify the cortical layer for each object output by Ilastik.

### Blood vessel segmentation

Individual blood vessels were manually segmented from clear brain images using a virtual reality image analysis software (Syglass). Blood vessels with diameters of approximately 20 µm were selected for segmentation, and 18-25 blood vessels were manually masked and segmented in each sample. Diameters were measured by drawing a line across the center cross-section of each blood vessel and averaging three measures taken from separate locations. Individually masked blood vessels were then realigned with their original image in Imaris, and distance transforms were calculated and exported for each blood vessel.

### Intensity & density calculation and binning (MATLAB r2023b)

MATLAB scripts were developed to calculate the intensity of tau staining, neuron density, and tau tangle density, along and away from the segmented blood vessels’ surfaces. First, tau data, segmented blood vessel distance transform images, and segmented cortical layer images were loaded simultaneously in one MATLAB script to align all images and export coordinates for data within 100 microns of the blood vessels’ surfaces. These coordinates contained data for each pixel in this range, with their X, Y, and Z positions; tau intensity; and cortical layer. To account for staining differences in each sample, tau intensity was normalized between samples using a piecewise linear normalization. The pixel intensity for background, autofluorescence, and AT8 positivity were recorded in each sample at 3 depths in the images’ z-stacks, with 10 measurements in each category per depth. Then, measurements were averaged for each category and linear functions between them were calculated to normalize the data on a singular dataspace.

To analyze tau staining intensity along the blood vessel, the exported coordinates were input into a script that calculates the pixel distance, in microns, along the surface of the blood vessel. This script calculated a centerline through the blood vessel, found the nearest position on the centerline to each tau pixel, and calculated that position’s distance from the start of the centerline^14^. The data was then exported as the original coordinates with their distance along the vessel appended. A similar script was used for the neuron and tangle density analysis, where the object coordinates obtained from Ilastik were input along with the intensity coordinates. This script calculated the distance along the vessel for each object within 30 microns of the vessel surface.

Finally, these data were binned into groups based on their distance along and away from the blood vessel. Immunolabeling intensity, neuron, and NFT data were grouped in 10-micron intervals along the vessel surfaces. For tau intensity, the mean intensity was calculated for each bin, and data within 3 microns from each vessel surface was used to determine the surface tau percentile^15^. For neurons and tangles, their density was calculated by measuring the number of objects within 30 microns from the vessel surface and comparing their quantity to the spatial volume of each bin.

## Results

### Three-dimensional histology reveals the accumulation of tau protein along blood vessels

Tissue was collected from the inferior temporal gyrus, a region associated with functional impairment and tau accumulation in AD^16^. Each block was sliced into 0.5-1 mm thick sections, then cleared and immunolabeled following an optimized protocol described previously^11^. AD and control human brain tissue samples immunolabeled for phospho-tau (AT8, recognizes tau phosphorylated at both S202 and T205), blood vessels (Glut1), neurons (HuD), and cell nuclei (DAPI, **Fig. 1A-B**)^17–20^.

**Figure 1.**
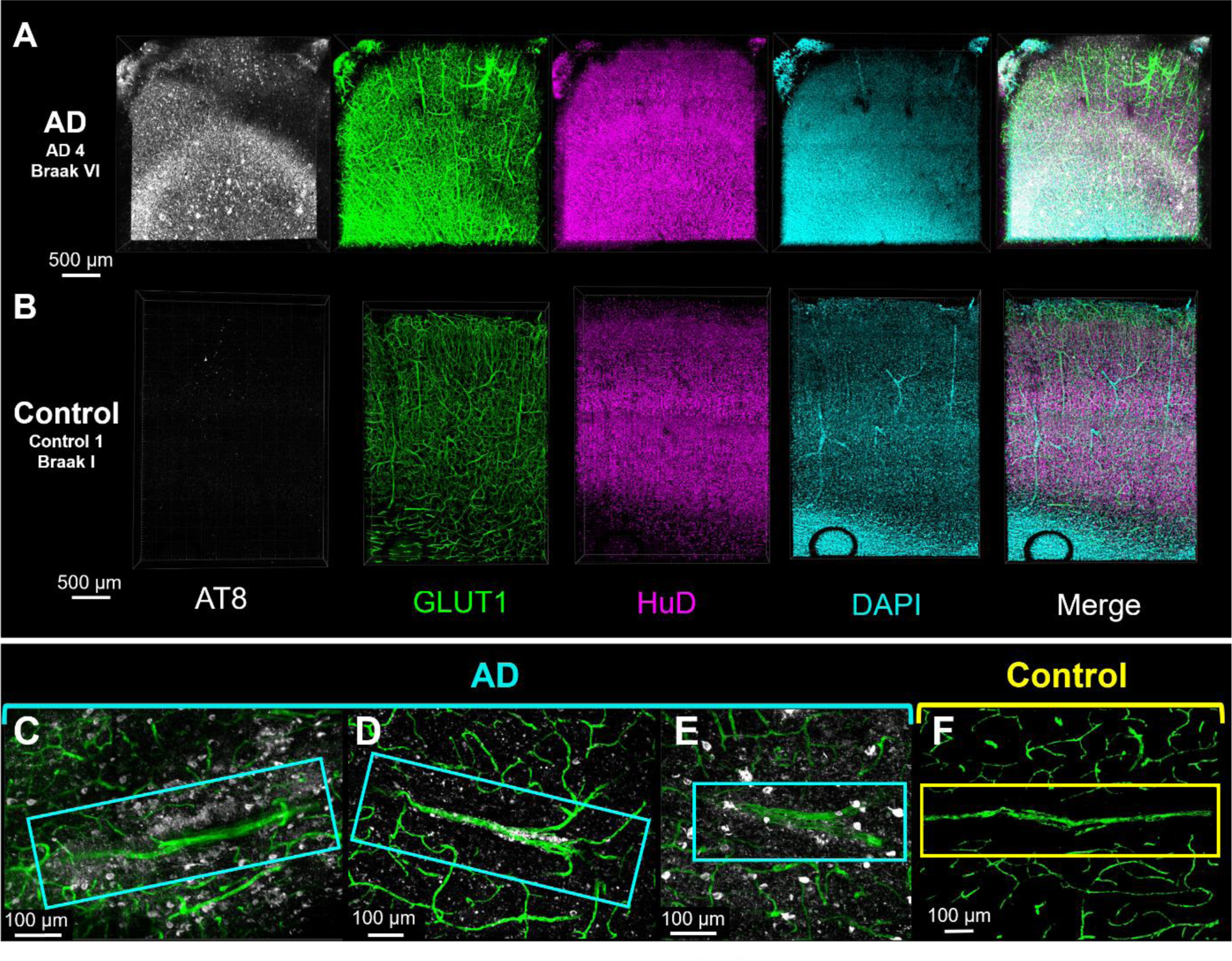
Three-dimensional histology reveals the presence of vascular-associated tau. Confocal fluorescence microscopy images showing raw data from the inferior temporal gyrus of an AD (A) and a control (B) donor. Images are immunolabeled for vasculature (GLUT1, green), neurons (HuD, magenta), nuclei (DAPI, blue), and AT8 tau (white). (**C-E**) Vascular tau accumulation in blood vessels AD donors compared with a **(F)** control donor. Images show staining for vasculature (green) and AT8 tau (white) in a 40 µm thick z-slice.

Visual inspection of vasculature reveals significant phospho-tau staining along the surface of some blood vessels (**Fig. 1C-E**). This phospho-tau staining is distinct from neurofibrillary tangles and shows a diffuse pattern that appears regionally along the length of select blood vessels in each sample. Control samples (Braak 0-I) did not have neurofibrillary tangles or phospho-tau + staining along blood vessels (**Fig. 1B, F**). In segments with vascular tau staining, staining also appears to extend away from the blood vessel surface and diminish as distance increases from the surface (**Fig. 1C, D**). Additionally, select blood vessels showed the accumulation of NFTs in addition to the diffuse staining on their surface (**Fig. 1E**). These observations suggest a spatial relationship between vasculature and phosphorylated tau accumulation in AD that is distinct from the accumulation of NFTs.

### Characteristics and cortical location of blood vessels with tau accumulation

To better define vascular tau, we established a protocol for isolating, quantifying, and classifying regions of tau accumulation on blood vessels. The virtual reality (VR) image analysis software Syglass was used to manually segment individual blood vessels from each sample by tracing masks along the surface of each blood vessel (**Fig 2A**). We found that VR tracing allowed us to more efficiently and accurately segment individual vessels compared to segmentation on 2D planes. Individual vessel masks were then realigned to the original image coordinates for the quantification of staining in other channels. In total, we segmented 18-25 blood vessels from n=6 AD and n=6 control donor brains. We also subdivided each image into its respective layers and found that the average tau intensity on the blood vessel surface—defined as the region within 3 microns of the vessel mask— was distributed across all cortical layers except for layer 6. This distribution pattern is distinct from the amount of tangles and neurons present in each cortical region (**Fig. 2 B, C**). Vascular tau was not observed along microvessels (<10µm in diameter), and the average diameter of measured vessels was 17 µm ± 4 (std. dev.). Vessels of similar size were selected for comparison in control tissues (**Fig. 2D**). Last, nearly all vessels with tau accumulation appeared to be arterioles, as indicated by co-labeling with smooth muscle actin (SMA, a marker of smooth muscle cells; **Fig. 2F**).

**Figure 2.**
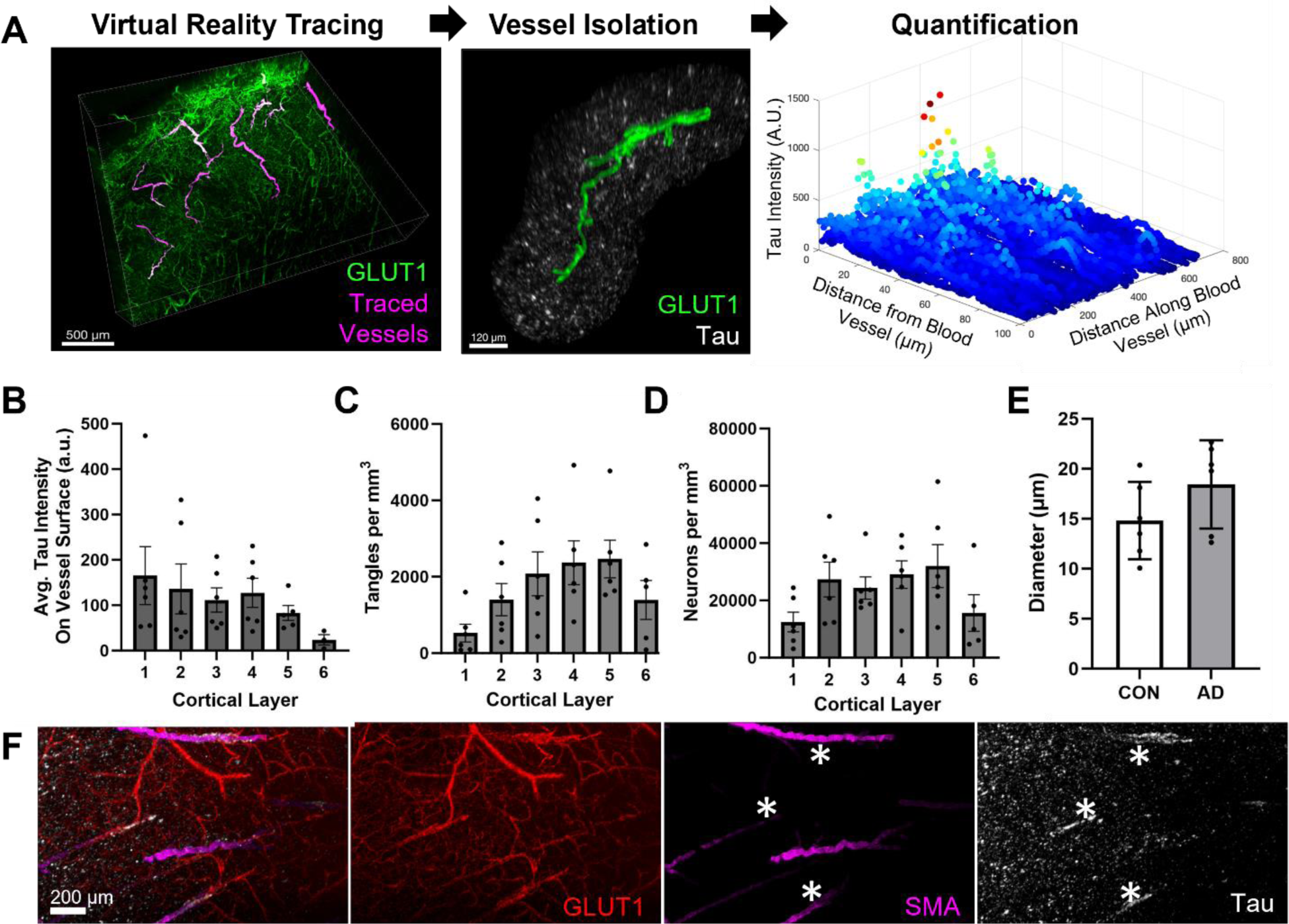
Examination of tau along individual blood vessels throughout the cortex. (**A**) An overall schema of the method used to quantify vascular tau. Blood vessels are first traced in virtual reality (magenta) and are shown overlying the original GLUT1-positive blood vessel imaging data (green). Tracing allows for the isolation of individual blood vessels and their surround, including tau pathology (white) ≤ 100 µm from the blood vessel surface (example is from AD 5 vessel 5). Subsequently, quantification of tau intensity along and away from the blood vessel surface was conducted. **(B)** Measures of the average tau intensity at the vessel surface (within 3 microns) per donor and cortical layer. **(C)** The average AT8-positive tangle density and (**D)** HuD-positive neuron density per cubic mm was also measured for each cortical layer. **(E)** The average diameter of vessels measured per donor. Dots represent individuals, bars represent means +/- SEM. **(F)** Separate tissue was labeled with antibodies to GLUT1, SMA, and tau show areas of tau accumulation on blood vessels that are also SMA-positive (indicated by asterisks).

### Frequency of vascular tau accumulation in inferior temporal gyrus

Next, we measured the intensity of tau labeling along the vascular surface. Each measured vessel had a segmented length that was continuous for roughly 300-2000 microns (**Fig. 3A**). Control samples consistently lacked tau accumulation along any blood vessels. There was also substantial diversity in the pattern of tau along blood vessels, both within and across AD samples. To simplify comparisons of tau accumulation across samples, we subdivided each 10-micron interval along a vessel surface into segments and assigned each segment a percentile based on the average phospho-tau staining intensity within that segment (**Fig 3B**). Regions of tau accumulation included stretches spanning small vascular lengths of <50 microns to >1000 microns and could appear continuous or patchy (**Fig. 3C-H**).

**Figure 3.**
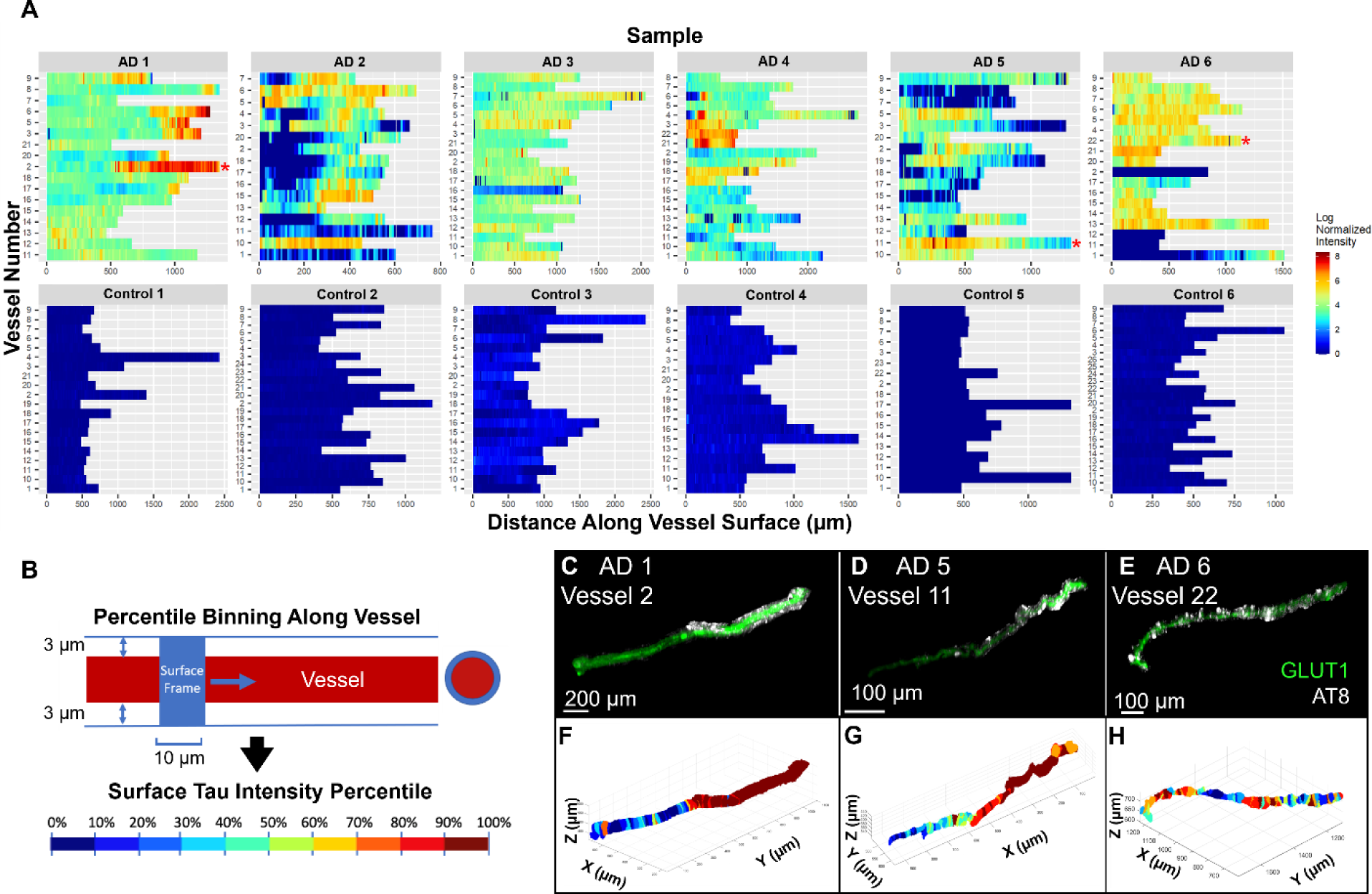
Mapping tau accumulation on blood vessels. **(A)** Heatmaps showing log normalized tau intensity within 3 microns from the surface of each segmented blood vessel (n=107 AD, n=127 control). Rows are individual vessels and columns are tau intensity measures along the vessel length. **(B)** Data is binned every 10 microns along the blood vessel’s surface and shows the mean intensity of each bin, normalized to the mean tau intensity of the whole image. Red asterisks highlight example vessels shown in panels C-H. **(B)** A schematic showing how data binning and surface tau measures were acquired. Each bin was then percentile ranked by AT8 tau staining intensity. **(C, D, E)** Example of isolated blood vessel (green) and tau labeling (white). **(F, G, H)** Corresponding maps of tau intensity along the vessel surface. Color corresponds to percentiles (deciles).

We hypothesized that one potential explanation for these observations could be that vessels occasionally travel through regions of high tau pathology. To rule out that the appearance of tau is incidental, we compared differences between groups of vascular segments within each percentile bin. Segments with the most vascular tau (top 90-100 percentiles) show substantial increase in tau intensity that decreases with distance from the vessel surface (**Fig. 4**). This indicates that tau is enriched near blood vessels compared to the surrounding tissue. By comparison, segments in the next decile (80-90^th^ percentile) show no substantial change in tau intensity with distance from the vessel surface, indicating no enrichment. Segments with less surface tau (80^th^ percentile and below) show the opposite trend—a decrease in tau intensity near the vessel surface (**Fig. 4**). Together, these data strongly support a relationship tau present near the vessel surface is distinct from that in the surrounding tissue.

**Figure 4.**
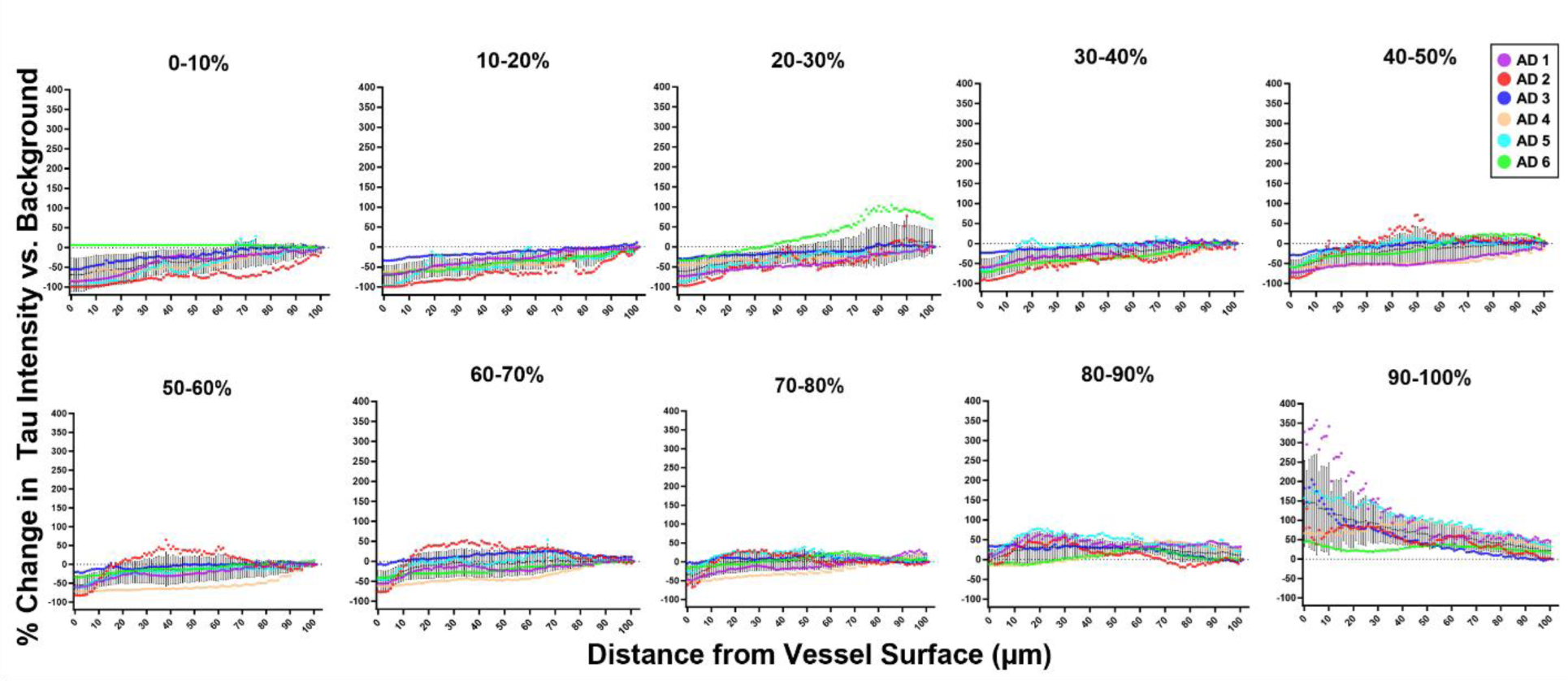
Tau intensity is related to distance from the blood vessel surface. Blood vessel segments were grouped according to surface tau intensity by percentiles (deciles) and the amount of tau immunolabeling is plotted by distance from blood vessel surface. Values for each donor (colored lines) are normalized to the average tau value in the whole image (background) such that a 0% change (grey dashed line) means the tau labeling intensity is no different than the average level of tau in the whole image.

### Composition of tau in blood vessels

Pathological tau is heavily post-translationally modified, so we additionally sought to understand what forms of tau are present in this vascular compartment by physically isolating blood vessels from the ITG of our AD and control samples and conducting WES assays (**Fig. 5A**). The assays revealed a significant increase total tau, the tau N-terminus (Tau13), and phospho-T181 and -T217 tau in the blood vessels of AD donors compared to controls (**Fig. 5B, C, E, G**). However, the levels of other forms of tau, S202, P231, and the tau C-terminus (Tau46), were not found in significantly higher levels in AD samples compared with controls (**Fig. 5D, F, H**). This indicates that enrichment of tau in vasculature is not an artifact of the AT8 antibody and that certain forms of tau are increased in this compartment in AD.

**Figure 5.**
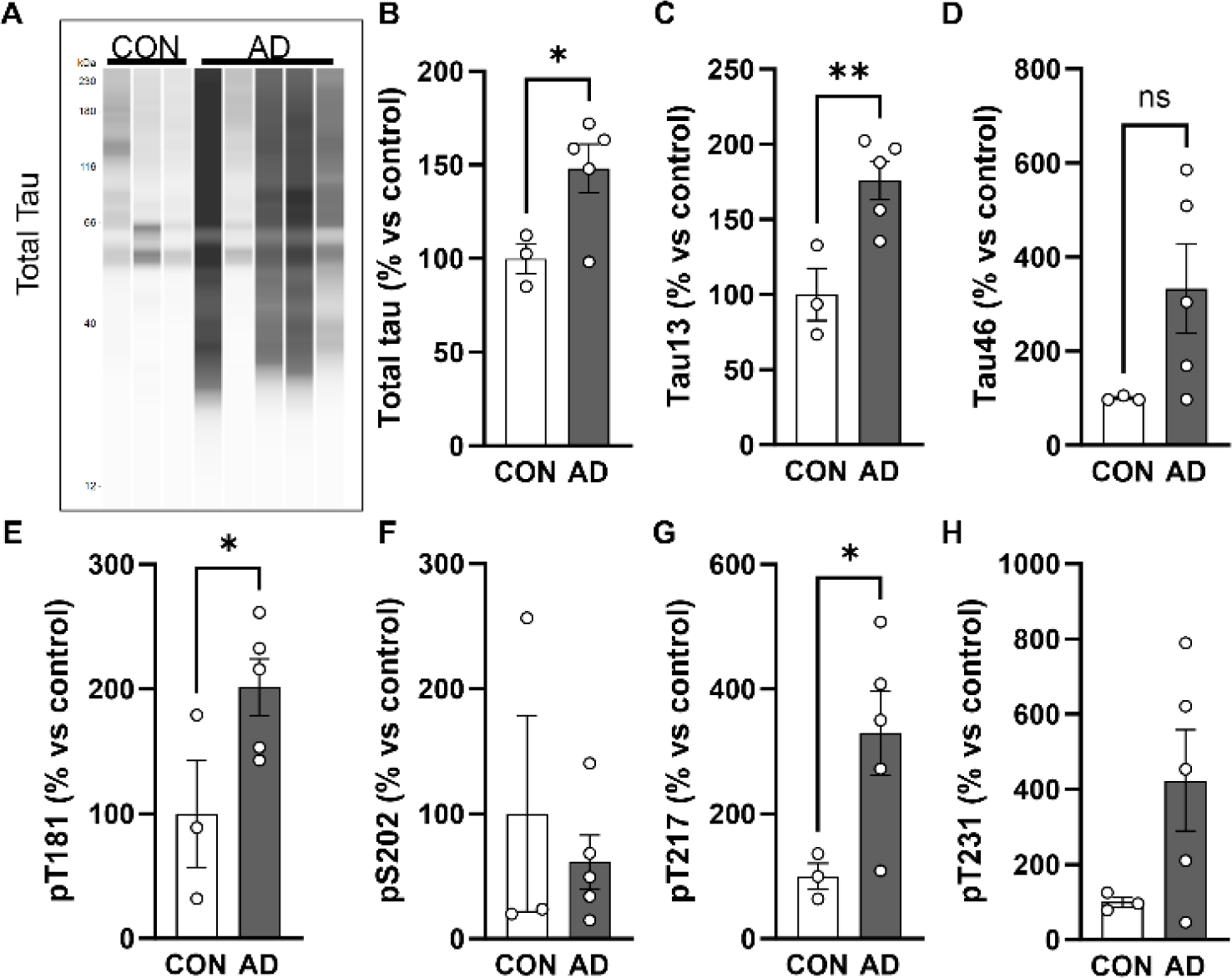
Post-translationally modified tau is enriched in AD blood vessels. **(A)** Isolation of blood vessels from the inf. temp. gyrus of n=3 control and n=5 AD brains shows that tau is enriched in vasculature and can be visualized with multiple antibodies including total tau = DAKO rabbit polyclonal. Quantification of total signal per lane for **(B)** total tau, **(C)** Tau13 n-terminal antibody, **(D)** Tau46 c-terminal antibody, **(E)** phospho-T181 tau, **(F)** phospho-S202 tau, **(G)** phospho-T217 tau and **(H)** phospho-T231 tau. All values normalized to the average of controls. One-tailed t-test *p<0.05, **p<0.01. Error bars = means +/- SEM.

### Relationship between vascular tau and NFT burden

Given the enrichment in vasculature for tau species known to contribute to the formation of NFTs, and an observation that tangles in AD tissues were frequently adjacent to tau-positive vessels (**Fig. 6A-D**), we next wanted to understand if areas of increased vascular tau were related to the local NFT burden. To do this, we segmented individual NFTs and neurons in our images (**Fig. 6E-G**). We then calculated the percent of neurons containing NFTs near blood vessels to determine the relationship between the amount of vascular surface tau and the likelihood of nearby neurons being NFT-positive. This quantification took place in the tissue immediately adjacent to the blood vessel--defined as a volume within 30 microns of the vessel surface.An ANOVA, correcting for repeated measures, shows a significant difference in the percent of neurons with NFTs that varies with vascular surface tau percentile (P value = 0.037, R^2^= 0.47; **Fig. 6H**). This indicates that as vascular surface tau increases, so too does the amount of nearby neurons with NFTs.

**Figure 6.**
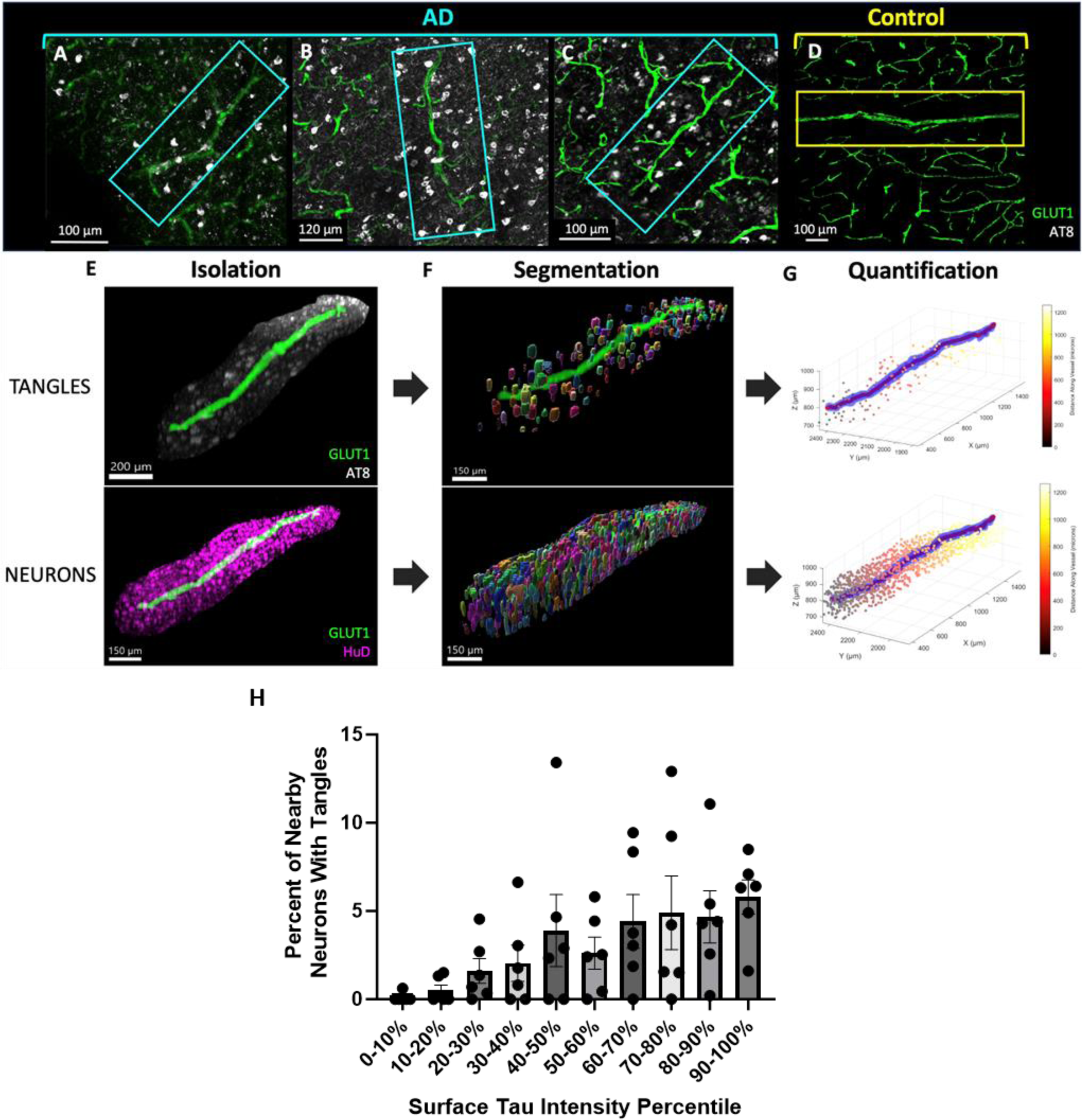
NFT and neuron density analysis. Examples of blood vessels showing NFT accumulation near the blood vessel surface in samples (A) AD 2, (B) AD 1, (C) AD6, and (D) control 3. (E) Isolated blood vessel with surrounding tau pathology (AT8, white) and surrounding neurons (HuD, magenta). (F) NFTs and neurons were identified and segmented using Ilastik. Visualization of segmentation masks were generated using Imaris and with a value of 1 μm to smooth the surfaces. (G) A custom MATLAB script was developed to calculate the distance of each segmented tangle and neuron from the surface of each blood vessel. Plots show the blood vessel (blue), its calculated centerline for object distance calculations (red), and objects colored according to their distance along the blood vessel. (H) Plot shows the percent of neurons with NFT in regions near (0-30 microns) blood vessels with varying amounts of surface tau. Repeated measures ANOVA P value = 0.037, R^2^= 0.47. Dots represent individuals, bars represent means +/- SEM.

## Discussion

Tau accumulation and NFT formation are closely associated with the clinical progression of Alzheimer’s disease^1^. Here, we report that the accumulation of tau along blood vessels is apparent in three-dimensional histology. These experiments indicate that pathological tau exhibits a close spatial relationship to vasculature in human Alzheimer’s disease brain. This provides evidence of an interaction between tau pathology and blood vessels, perhaps similar to CAA. Such an interaction reinforces previous studies conducted in mouse models of tau pathology which found changes in cerebral microvessels^10, 21, 22^. These results also provide further evidence for the existence of a vascular clearance pathway for tau pathology, which may be disrupted in regions with increased levels of tau accumulation.

Recent studies have investigated the role of vasculature in tau clearance at larger, systemic scales in mouse models of AD, finding that brain vasculature and the associated glymphatic system regulate clearance of tau pathology^6, 7, 23^. These conclusions have been supported in human magnetic resonance imaging, where perivascular spaces were found to be associated with tau pathophysiology in early AD^24^. Our study investigated the association of tau and vasculature at smaller, single-cell spatial scales in human AD donors to determine if patterns of tau accumulation are consistent with the observations of these previous studies. Indeed, large-volume images we obtained of the inferior temporal gyrus indicate that tau accumulates along vascular segments in AD, suggesting that the clearance of tau pathology by vasculature may be dysfunctional in this area. Specifically, we found that some blood vessels exhibit regions with higher levels of tau near the blood vessel surface than the surrounding tissue--indicating that tau-positivity is not simply occurring by chance. Additionally, in areas with low levels of vascular tau, tau intensity is lower near the vessel surface compared to its surroundings, suggesting functional vascular clearance. Altogether this supports the idea that vessels are important for tau clearance.

Overall, the deposition of tau on blood vessels is similar to CAA. Tau-positive vessels are primarily arterioles. However, leptomeningeal vessels were not observed to be sites of tau deposition, with areas of tau-positive vessels being distributed throughout cortical layers I-V. By comparison, CAA type 2-affected vessels are frequently found in the leptomeninges and arterioles in superficial cortical layers ^25, 26^. Tau-positivity also did not appear to overlap with dyshoric capillary changes, though cases with CAA type 1 were not included in this study^27^. Prior studies have reported that neuritic dystrophies are increased near vessels that accumulate CAA^28, 29^. While we did observe dystrophic neurites around some blood vessels, neither these nor the additional diffuse staining along the length of vessels appeared to be directly related to CAA. However, we cannot rule out that some tau-positive vessels may also be affected by CAA at other locations along their length. Other CAA-related features that we did not observe in tau positive segments included vessels with a “double-barrel” appearance or the presence of microhemorrhages, though direct examination of CAA-positive tissues would help to confirm these observations.

Furthermore, we performed protein assays that found blood vessels in AD patients contain higher levels of phospho-tau species within the blood vessel walls themselves compared to controls. In particular, we observed increased phospho-T217. Phospho-T217 tau is also a sensitive blood-based biomarker for AD and a form of phosphorylated tau that has been found to accumulate in AD and drive the hyperphosphorylation and fibrillization of wild-type tau^30, 31^. This suggests that blood vessels harbor aggregate-prone species of tau, which was further supported by analysis showing an association between vascular tau and local NFT density. These data are in line previous reports that observed a greater incidence of tau labeling near vessels with increasing Braak stage and our own prior work showing that isolated blood vessels from AD brain are enriched for tau species capable of seeding new aggregates^10, 32^. Together, these observations suggest that impaired vascular clearance of tau may contribute to the progression of AD pathology.

In addition to the experimental results of our study, we also present a new methodology for characterizing disease pathology relative to anatomical structures. Until recently, it has not been possible to image large tissue volumes with cellular and sub-cellular resolution, but new imaging methods, such as confocal and light sheet fluorescence microscopes, coupled with tissue clearing technology have now enabled this. However, many of the current, most widely used quantification tools face challenges analyzing these images, as they were primarily designed for traditional, two-dimensional analysis^33,34^. This study presents a method utilizing emerging machine learning and virtual reality tracing software to identify objects throughout large images, while accounting for differential staining and object characteristics throughout the image volume, a challenge that traditional simple thresholding and rolling-ball filtering methods do not account for. This is a significant development, because it allows for the alignment of pathology, brain structures, and original imagery to investigate spatial relationships across large regions, while maintaining cellular or subcellular resolution.

While these data indicated that tau is enriched at points along blood vessels compared to the surrounding tissue, additional characterization of tau-enriched vessels is needed to better understand the cause and consequence of this buildup on blood vessels. For example, studies have implicated aquaporin 4 (AQP4) in the clearance of tau pathology; thus, AQP4 provides a possible target for studies looking to determine the precise cause of vascular tau^35^. If specific transporters or tau-interacting proteins can be identified, they may provide a new target for therapeutics designed to remove tau pathology. Further, while nearly all vessels examined in AD inferior temporal gyrus exhibited regions of enhanced tau accumulation, whether these findings can be extended to other brain areas, including those where NFTs are relatively scant such as the visual cortex and cerebellum, could widen our understanding of this phenotype.

In summary, this study provides new evidence of brain vasculature’s role in the progression of AD and distribution of pathology. Perhaps most notably, our results indicate that tau deposits around vasculature with characteristics similar to amyloid beta in CAA. Additionally, this work provides further support for the role of vasculature in mediating tau clearance. Further investigation of how this disrupts vascular functions including specific transporter mechanisms in endothelial cells, may help uncover new methods to modify tau burden in the brain via vascular clearance.

## Funding and Acknowledgements

This work was supported by a grant from the Alzheimer’s Association (23AARG-1029355; REB), by the NIH NIA R00AG068602 (TJZ) and R00AG061259 (REB), the Harrison Gardner Jr Innovation Award (BTH, TJZ), the Jack Satter Foundation (BTH, REB, TJZ), and a Predoctoral Fellowship FPU18/00630 from the Spanish Ministry of Science, Innovation, and Universities (ES). We additionally thank the donors and their families who have contributed to the Massachusetts Alzheimer’s Disease Research Center, which is supported by the NIH NIA P30AG062421. We would also like to thank the Harvard Center for Biological Imaging including Douglas Richardson for the use of their imaging facility and helpful conversations.

